# Distinct dermal fibroblasts direct mechano-chemical signaling to the epidermis during pregnancy

**DOI:** 10.1101/2025.09.11.675503

**Authors:** Yoshihiko Kobayashi, Kazunori Sunadome, Koichiro Maki, Hiroki Fukunaga, Hitomi Matsubara, Sahomi Ohkubo, Ritsuko Maki, Maki Yoshikawa, Aleksandra Tata, Purushothama Rao Tata, Ken-ichi Matsumoto, Taiji Adachi, Mitsuhiro Iwaki, Takuya Yamamoto, Fumiko Toyoshima

**Author notes:** Corresponding Authors. Email: Fumiko Toyoshima, Takuya Yamamoto, Mitsuhiro Iwaki.

## Abstract

Dermal fibroblasts alter extracellular matrix (ECM) and tissue mechanics dynamically in wound healing and fibrosis, however, are understudied in healthy skin remodeling across life stages. Here, by conducting spatio-temporal transcriptomics, we identified *Cdh4*^*+*^ dermal fibroblast subpopulation that remodels ECM and converts mechano-to-chemical signals to promote epidermal stem cell proliferation in expanding abdominal skin of pregnant mice. Mechanistically, *Cdh4*^*+*^ fibroblasts produce a pregnancy-responsive matrisome, resulting in denser and stiffer dermal fibrils. These fibroblasts then sense the stiffened substrate and convert the mechanical input into a chemical output via the YAP1-TGFβ2 axis, leading to the proliferation of epidermal stem cells via TGFβ receptor in the expanding skin. Thus, *Cdh4*^*+*^ fibroblasts fine-tune the dermal mechanofield and act as a mechanical-to-chemical signal conversion hub in healthy skin expansion.

## Main text

Tissue stiffness influences stem cell behaviors including self-renewal, differentiation and cell-fate decision, contributing to the remodeling of tissue architecture (*1*–*3*). The epidermal stem cells in the skin change their properties depending on the stiffness of the basement membrane to which they are attached, resulting in tumor formation and age-dependent stem cell exhaustion (*4, 5*). Recent studies have emphasized the importance of dermal stiffness underlying the basement membrane in relation to the compromised behavior of epidermal stem cells in aging and pathological conditions (*6, 7*). However, the relevance of dermal mechanical properties to epidermal stem cell dynamics in healthy skin remodeling remains poorly understood.

During pregnancy, the skin on the abdomen expands to accommodate the growing fetus. A failure of the skin expansion can cause striae gravidarum (SG; known as stretch marks), that is suggested by the observation that the people with Ehlers-Danlos syndrome (EDS), whose skin shows over flexibility and fragility, have no SG after pregnancy (*8*). Matrisome gene expression pattern alters in striae fibroblasts (*9*), suggesting that unsuccessful remodeling of dermal extracellular matrix (ECM) leads to stretch marks. In mice, the emergence of highly proliferative interfollicular epidermal stem cells (IFESCs) in the abdominal skin during pregnancy is regulated by blood vessels and signals from dermis (*10, 11*), however, the insights of dermal ECM remodeling is limited.

The dermis, a rich reservoir of fibrillar collagen and other ECM components, provides crucial mechanical cues to the skin tissue (*12*). As the major source of ECM, fibroblasts are highly plastic cells that adapt to the micro-environment of their anatomical region and physiological changes of the body (*13*–*15*). Beyond this structural role, fibroblasts are increasingly recognized as central chemical signaling hubs, secreting a wide array of growth factors and cytokines that orchestrate the behavior of surrounding cells (*16, 17*).

Here, we identify a distinct subpopulation of dermal fibroblasts that alters the dermal mechanical cues of pregnant mice by secreting pregnancy-responsive ECM. This fibroblast subpopulation also functions as a central signaling hub, converting mechanical cues of stiffening ECM into chemical signals that drive the proliferation of IFESCs during pregnancy. Our findings reveal a mechanism of dermal stiffness-dependent regulation of epidermal stem cells in healthy skin expansion orchestrated by a specialized fibroblast signaling hub.

### A distinct dermal fibroblast subpopulation exhibits dynamic ECM gene expression during pregnancy

To explore the transcriptomic landscape and its transition of the dermis during healthy tissue remodeling at single-cell level, spatio-temporal transcriptomic analysis of pregnant mouse abdominal skin (4-, 9- and 17-days post coitum (dpc)) was performed by combining single-cell RNA-seq (scRNA-seq) and spatial transcriptomics (Fig. 1A). We designed the analysis with using the cells after removing epidermis to investigate the cellular responses in dermis. With the dermis enrichment cell isolation and 10x Genomics Chromium scRNA-seq, total 12,862 dermal and 2,059 epidermal cells were utilized for finalized analysis after filtering low quality cells (fig. S1, A to C). Sub-clustering of the fibroblast population revealed at least five subpopulations of fibroblasts, including *Cdh4*^+^, *Coch*^+^, *Plac8*^+^, *Slc6a6*^+^ and *Sox9*^+^ subpopulations, exist through gestation and in the non-pregnant control (Fig. 1B and fig. S1D). To resolve the spatial distribution of these fibroblast populations, we performed hybridization-based in situ sequencing (HybISS) (*18*) with several optimization (fig. S2A). The primary modification was omission of the reverse transcription step, which improved detection efficiency in skin tissue. This optimized protocol (hereafter RNA-HybISS) enabled us to detect markers for multiple layers of mouse skin tissue (fig. S2B). Using a panel of 66 marker genes for the five fibroblast subpopulations (Fig.1C) in combination with probabilistic cell typing by in situ sequencing (pciSeq) (*19*), we identified *Cdh4*^+^ fibroblasts as the major subpopulation in the dermis. *Plac8*^+^ fibroblasts, meanwhile, were dominant in the hypodermal layer (Fig. 1, D, E, fig. S2, C and D). Reactome pathway enrichment analysis revealed that *Cdh4*^+^ fibroblasts preferentially express genes associated with ECM organization compared to other fibroblast subpopulations (Fig. 1F). Principle component analysis restricted to matrisome-related genes demonstrated that the expression patterns in *Cdh4*^+^ fibroblasts were substantially altered in mid-to-late pregnancy (Fig. 1G). A few genes, including *Tnxb, Col4a1* and *Thbs2*, were upregulated at 4 dpc, whereas most genes, including *Col1a1, Col3a1, Lox, Eln* and *Sparc*, began to increase their expression during mid-to-late pregnancy (Fig. 1H), implying ECM reorganization starts in early pregnancy and continue through gestation. Taken together, *Cdh4*^+^ fibroblasts are likely key modulators of dermal ECM alteration during gestation.

**Fig. 1.**
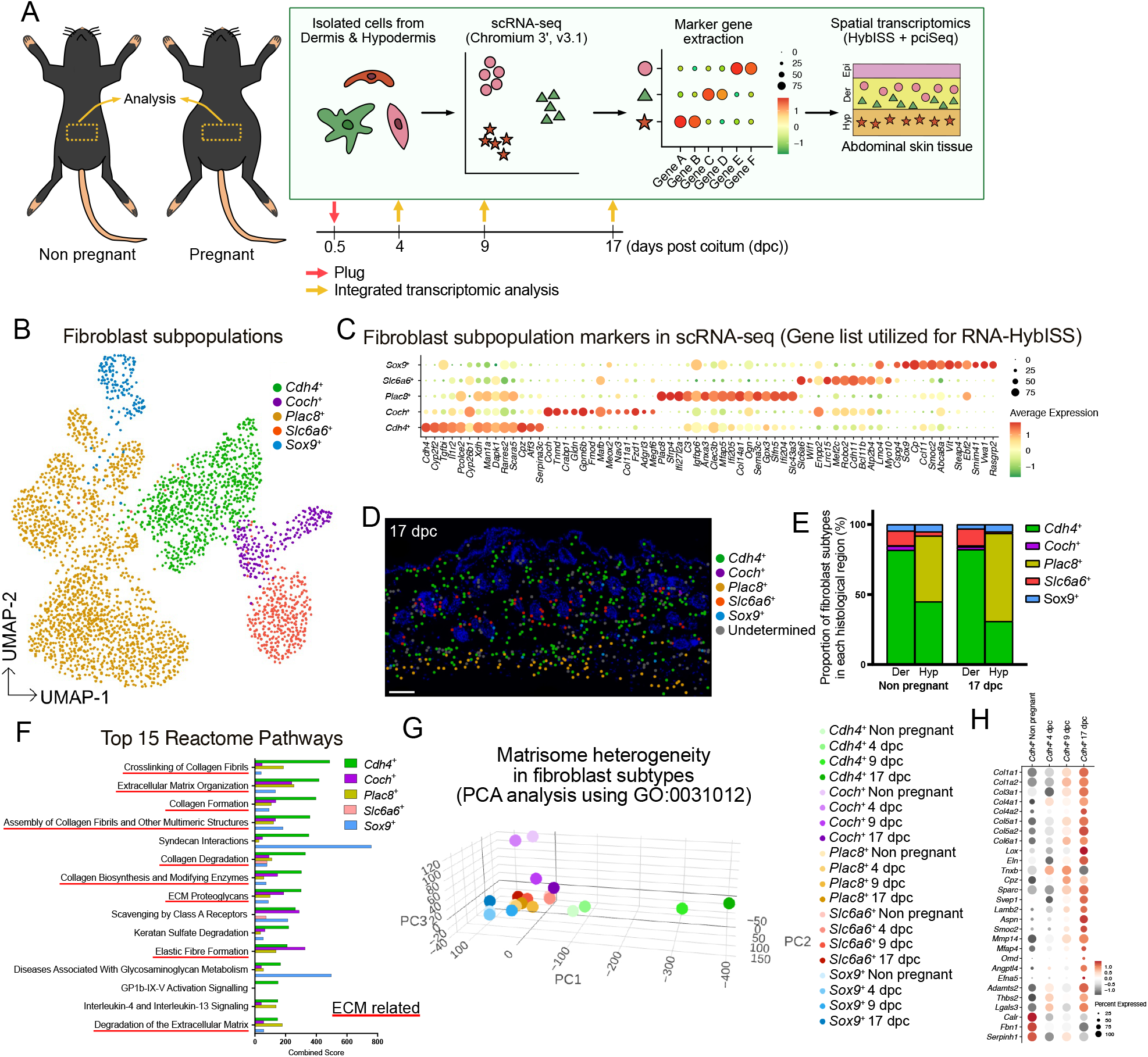
A distinct dermal fibroblast subpopulation exhibits dynamic ECM gene expression during pregnancy. (**A**) Schematic of integrated transcriptomic analysis of abdominal skin obtained from pregnant (4, 9, 17 days post coitum (dpc)) and non-pregnant mice. Epi, epidermis; Der, dermis; Hyp, hypodermis. (**B**) UMAP visualization of scRNA-seq of the dermal and hypodermal fibroblasts. (**C**) Dot plot showing markers for distinct five populations of the fibroblasts. The markers utilized for RNA-HybISS are shown. (**D**) Pci-Seq determination of the distribution of fibroblast subpopulations. Fibroblast subpopulations are represented by circles of the indicated colors. Dark blue indicates Hoechst staining of nuclei in non-fibroblast cell types. (**E**) Proportion of fibroblast subpopulations in Dermis (Der) and Hypodermis (Hyp) in non-pregnant and 17 dpc abdominal dermis determined by pci-Seq. (**F**) Reactome pathways enriched in *Cdh4*^+^ fibroblasts. Red underlines indicate ECM-related ontologies. (**G**) PCA plot of fibroblast subpopulations calculated based on genes listed in GO:0031012. (**H**) Dot plot showing matrisome genes regulated in *Cdh4*^*+*^ fibroblast during pregnancy. Scale bar: 100 μm (**D**).

### ECM remodeling induces stiffened dermal fibrils during pregnancy

To detect the dermal architecture and its transition during pregnancy, we performed Quantitative Imaging^™^ analysis of the abdominal dermis using atomic force microscope (AFM) (Fig. 2A). Quantitative Imaging^™^ analysis showed that dermal fibrils tend to become denser during pregnancy (Fig. 2B). In addition, the net stiffness of the dermal fibrous structure, as measured by AFM, increased significantly at 9 dpc compared to non-pregnant controls (Fig. 2C). Tenascin XB (*Tnxb*), a gene up-regulated in early-to-mid gestation in *Cdh4*^+^ fibroblasts (Fig. 1H), has been reported to promote collagen fibrillogenesis and increase the integrity of connective tissues (*20, 21*). In addition, the deficiency of *Tnxb* gene is known to induce classical-like EDS (clEDS1) that causes hyperextensible skin (*21, 22*). Quantitative Imaging^™^ analysis indicated that the dermal fibrils of constitutive *Tnxb* knockout (KO) mice at 17 dpc tend to become thinner (Fig. 2, D, E and fig. S3A), and lower net stiffness (Fig. 2F) than that of wildtype. These results indicate that TNXB-dependent ECM remodeling causes stiffening of dermal fibrils during pregnancy.

**Fig. 2.**
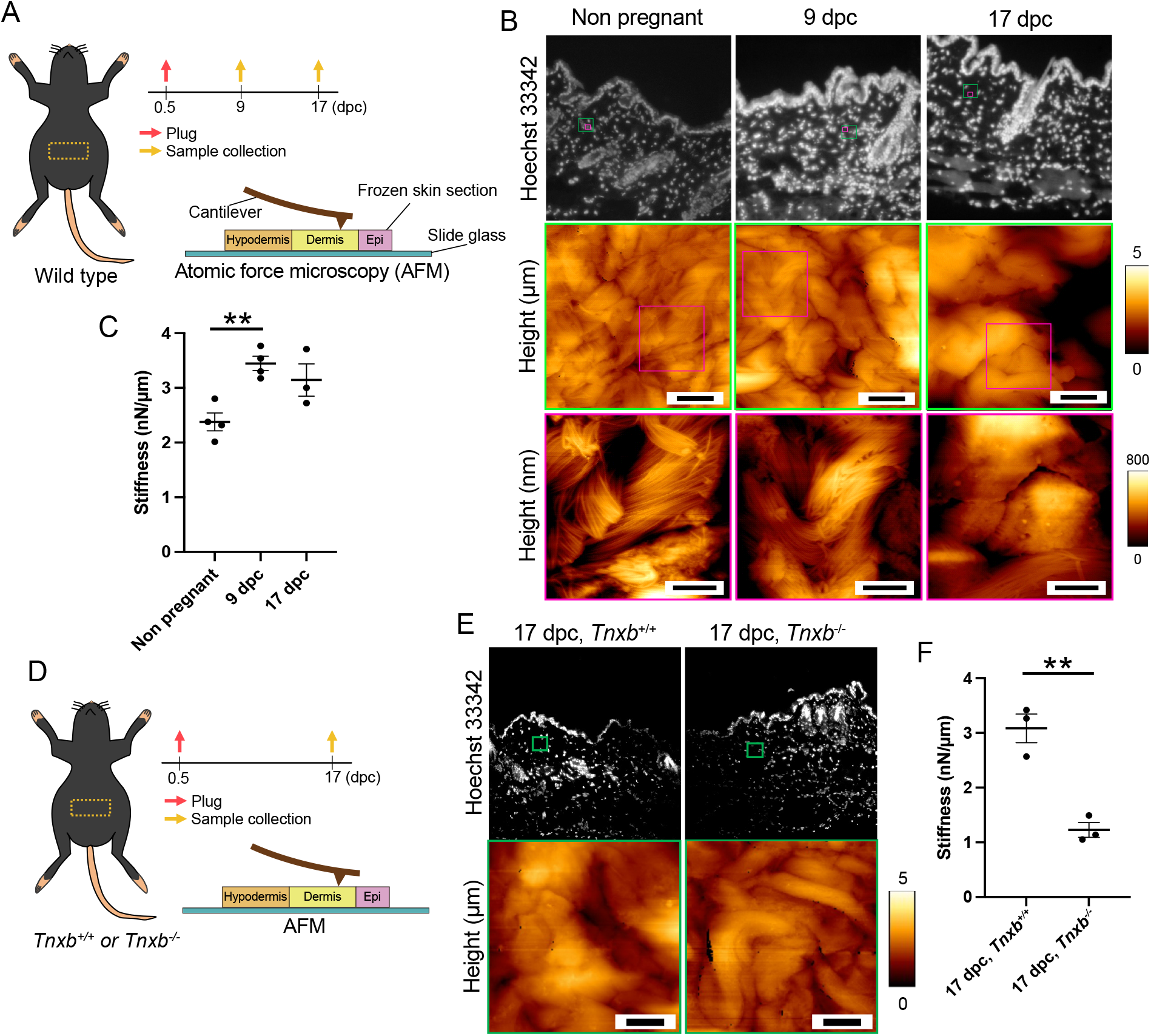
ECM remodeling induces stiffened dermal fibrils during pregnancy. (**A**) Schematic of atomic force microscopic (AFM) investigation of mouse abdominal skin tissues during pregnancy. (**B**) Fluorescent microscopic images of nuclei stained by Hoechst 33342 (top); heatmap of the tissue section height (middle and bottom). (**C**) Quantification of the net stiffness. Data are presented as the mean ± s.e.m. from three-to-four biological replicates. \*\**P* < 0.01, evaluated with one-way analysis of variance (ANOVA) with Tukey’s multiple comparison. (**D**) Schematic of AFM investigation of *Tnxb* KO mouse abdominal dermis during pregnancy. (**E**) Heatmap of the tissue section height. (**F**) Quantification of the net stiffness. Data are presented as the mean ± s.e.m. from three biological replicates. \*\**P* < 0.01, evaluated with two-tailed unpaired Student’s t-test. Scale bars: 5 μm (B, middle and E) and 2 μm (B, bottom).

### The stiffness that mimics the dermis in pregnancy activates mechanical signaling in fibroblasts

To investigate the functional consequences of the abdominal dermal stiffening observed during gestation, we examined the responses of dermal fibroblasts to hydrogel that can be adjusted for stiffness using newly established isolation and culture methods for primary mouse abdominal dermal fibroblasts (Fig. 3A, fig. S4, A and B). The stiffness of the hydrogel was tuned to mimic that of the abdominal dermis of a non-pregnant or pregnant mice at 9 dpc using AFM analysis (Fig. 3B). To directly measure the tension between fibroblasts and the hydrogel, we developed a DNA-based digital tension sensor (Fig. 3C). This sensor links a single integrin molecule on the cell membrane to the hydrogel, and the DNA strand modified with cholesterol and Cy3 dye dissociates from its complementary strand when the applied force exceeds the threshold (30 pN) (*23*). The released cholesterol strand anchors to the adjacent cell membrane (Fig. 3C), enabling the cells under larger and sustained forces to exhibit stronger Cy3 fluorescence signals (*24*). This technology allows cells to be re-seeded onto glass-bottom dishes for quantitative fluorescence imaging, which is advantageous for our culture system that is otherwise challenging to analyze directly due to the thickness and lack of surface flatness of the hydrogel. (Fig. 3C, fig. S5A). The results showed that the stiffness that mimics the dermis in pregnancy induced higher tension compared to the non-pregnant stiffness (Fig. 3D to F). To elucidate the downstream pathways following dermal fibroblasts sense the tension between themselves and matrix, we performed pathway enrichment analysis using the scRNA-seq data. Intriguingly, the pathway analyses reveal an enrichment of focal adhesion signaling (fig. S6A) and suggests the activation of Yes-associated protein 1 (YAP1), a mechano-transducer, in dermal fibroblasts during pregnancy (fig. S6B). In further *in vitro* analysis, the stiffened hydrogel that mimicked the dermis in pregnancy induced significantly higher nuclear translocation of YAP1 than that with the non-pregnant stiffness (Fig. 3, G and H). These data demonstrate that the dermal ECM stiffening observed in pregnancy is sufficient to activate mechanosignaling in resident fibroblasts.

**Fig. 3.**
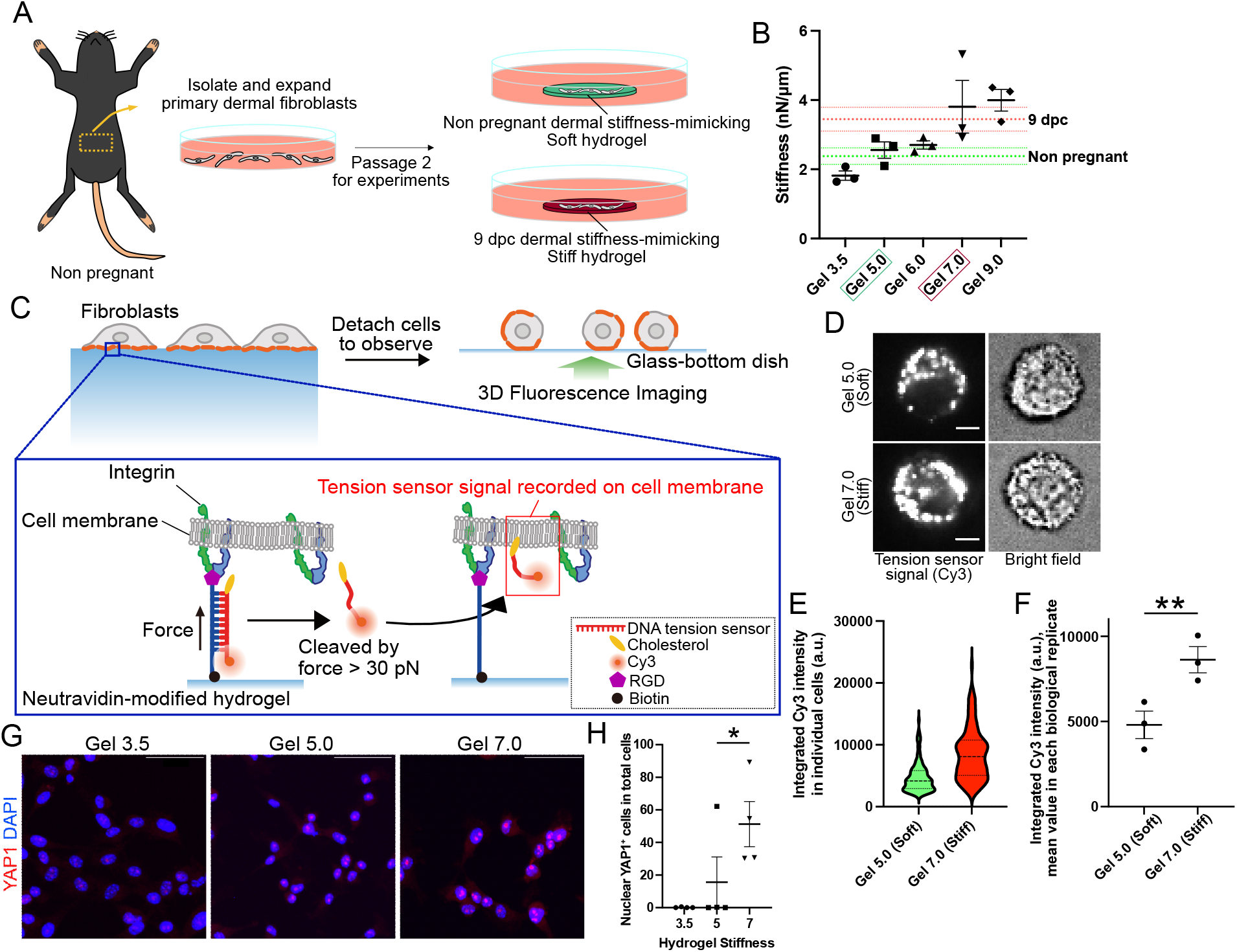
The stiffness that mimics the dermis in pregnancy activates mechanical signaling in fibroblasts. (**A**) Schematic of isolation and culture of mouse abdominal dermal fibroblasts on the stiffness-tuned hydrogel. (**B**) The net stiffness of hydrogel measured by AFM. Green and red dotted lines show net stiffness of dermis from non-pregnant and pregnant (9 dpc) mice, respectively, measured by AFM (±10%) (shown in Fig. 2C). Data are presented as the mean ± s.e.m. from three technical replicates. (**C**) Schematic of fibroblast culture on stiffness-tuned hydrogel with digital tension sensor. When tension exceeds 30 pN, Cy3-labeled ssDNA is released and incorporated into the cell membrane, enabling fluorescence-based readout. While both tensile and compressive forces act on the sensor, it responds specifically to tensile forces > 30 pN. Cells are then detached and re-seeded to glass-bottom dishes for accurate imaging. (**D**) Cy3 tension sensor signal of fibroblasts cultured on Gel 5.0 and Gel 7.0 coated with digital tension sensor for 6 h and re-seeded to glass-bottom dishes. (**E**) Integrated Cy3 intensity in individual cells. More than 34 cells for each replicate were analyzed. (**F**) Mean of (**E**) per biological replicates. Data are presented as the mean ± s.e.m. from three biological replicates. \*\**P* < 0.01, evaluated with two-tailed paired Student’s t-test. (**G**) Immunofluorescent images for YAP1 (red) in cultured abdominal dermal fibroblasts on stiffness-tuned hydrogel for 24 h. (**H**) Quantification of fibroblasts with YAP1 in their nuclei shown in (**E**). Data are presented as the mean ± s.e.m. from four biological replicates. \**P* < 0.05, evaluated with one-way ANOVA with Tukey’s multiple comparison. Scale bars: 5 μm (**D**) and 50 μm (**F**).

### YAP1 activated in *Cdh4*^+^ fibroblasts controls proliferation of IFESCs

We next sought to explore YAP1 activity and functions *in vivo*. Analysis of our scRNA-seq data revealed high expression of YAP target signature genes in several fibroblast subpopulations, including *Cdh4*^*+*^ fibroblasts (Fig. 4A). Furthermore, immunostaining showed that a greater population of dermal fibroblasts exhibited nuclear YAP1 localization during pregnancy compared to the non-pregnant dermis (Fig. 4, B to D). Consistent with the decreased stiffness (see Fig. 2F), the dermis of *Tnxb* KO mice showed a significantly lower percentage of fibroblasts exhibiting nuclear YAP1 (Fig. 4, E to G). These results indicate that dermal stiffening induces YAP activation in fibroblasts during pregnancy. The rapid expansion of abdominal skin in late pregnancy is known to require robust IFESC proliferation in mice (*10*), however, interactions between the mechanosignaling in dermis and IFESC proliferation remain unclear. To determine the contribution of the observed mechanosignaling pathway to the skin expansion, we performed double conditional KO (cKO) of *Yap1* and its orthologue *Wwtr1* (a.k.a. TAZ) in two types of fibroblasts: those expressing the pan-fibroblast marker *Pdgfra*, and those expressing *Cdh4* (Fig. 4H). The *Yap1/Wwtr1* double cKO in both types of fibroblasts significantly reduced the proliferation of abdominal IFESCs during pregnancy (Fig. 4, I, J), demonstrating that activation of YAP1 in *Cdh4*^+^ fibroblasts is necessary for IFESC proliferation. Taken together, *Cdh4*^+^ fibroblasts can act as a central hub that produces pregnancy-responsive ECM and converts mechanical cues from the ECM into biochemical signals, activating the stem cell functions of the epidermis and leading to skin expansion.

**Fig. 4.**
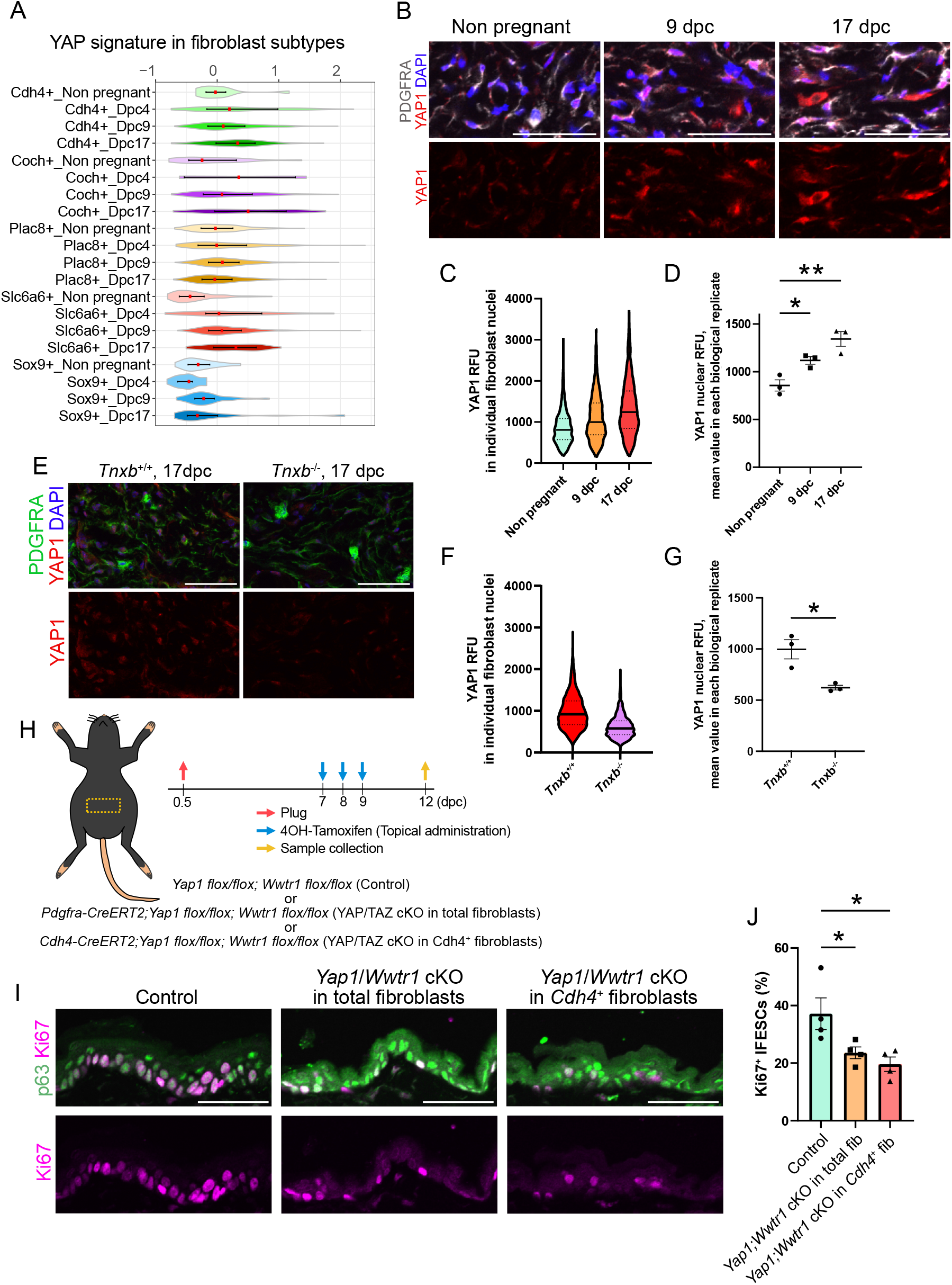
YAP1 activated in *Cdh4*^+^ fibroblasts controls proliferation of IFESCs. (**A**) Violin plot showing YAP signature expression levels in five fibroblast subpopulations in non-pregnant and pregnant abdominal dermis detected by scRNA-seq. (**B**) Immunostaining for YAP1 (red) and PDGFRA (gray) in abdominal dermis. (**C**) Violin plot showing YAP1 RFU in individual fibroblast nuclei shown in (**B**). (**D**) Mean of (**C**) per biological replicates. Data are presented as the mean ± s.e.m. from three biological replicates. \**P* < 0.05, \*\**P* < 0.01, evaluated with one-way ANOVA with Dunnet’s multiple comparison. (**E**) Immunostaining for YAP1 (red) and PDGFRA (green) in *Tnxb* KO mouse abdominal dermis at 17dpc. (**F**) Violin plot showing YAP1 RFU in individual fibroblast nuclei shown in (**E**). (**G**) Mean of (**F**) per biological replicates. Data are presented as the mean ± s.e.m. from three biological replicates. \**P* < 0.05, evaluated with one-way ANOVA with Dunnet’s multiple comparison. (**H**) Schematic of *Yap1*/*Wwtr1* double cKO experiments. (**I**) Immunostaining for Ki67 (magenta) and p63 (green) in abdominal skin of the *Yap1*/*Wwtr1* double cKO in total fibroblasts (middle) and *Cdh4*-expressing fibroblasts (right) and CreERT2 negative control (left). (**J**) Quantification of (**I**). Data are presented as the mean ± s.e.m. from four biological replicates. \**P* < 0.05, evaluated with one-way ANOVA with Dunnet’s multiple comparison. Scale bars: 50 μm (B, E and I).

### *Cdh4*^+^ fibroblasts induce IFESC proliferation through TGFβ signaling

To identify the paracrine factors targeted by YAP1 and released from *Cdh4*^+^ fibroblasts that promote IFESC proliferation, we performed scRNA-seq on the abdominal dermis of pregnant *Yap1*/*Wwtr1* double cKO mice (Fig. 5, A, B, fig. S7, A and B). Differential NicheNet analysis predicted that multiple genes encoding secreted proteins, including transforming growth factor beta 2 (*Tgfb2*), are the downstream factors of YAP/TAZ in *Cdh4*^+^ fibroblasts and potentially bind to the receptors expressed on IFESCs (Fig. 5C). Notably, *Tgfb2* was upregulated in *Cdh4*^+^ fibroblasts during pregnancy (Fig. 5D). ChIP-Atlas database (*25*) indicates transcriptional factors TEAD1 and TEAD4 and their co-factor YAP1 can bind to around the transcription start site of the *Tgfb2* locus (fig. S7C). Furthermore, gene set enrichment analysis (GSEA) of publicly available RNA-seq dataset of IFESCs (*11*) confirmed significant activation of the TGFβ signaling pathway in IFESCs during pregnancy compared to non-pregnant control (Fig. 5E). Consistent with this pathway activation, cKO of the TGFβ receptor 2 (*Tgfbr2*) in *Krt14*-expressing epidermal stem cells resulted in significantly less proliferative IFESCs during pregnancy (Fig. 5, F to H). Collectively, our study delineates that *Cdh4*^+^ fibroblasts act as a mechano-to-chemical signaling hub via a YAP1-TGFβ2 axis, governing a fundamental mechanism of physiological tissue remodeling.

**Fig. 5.**
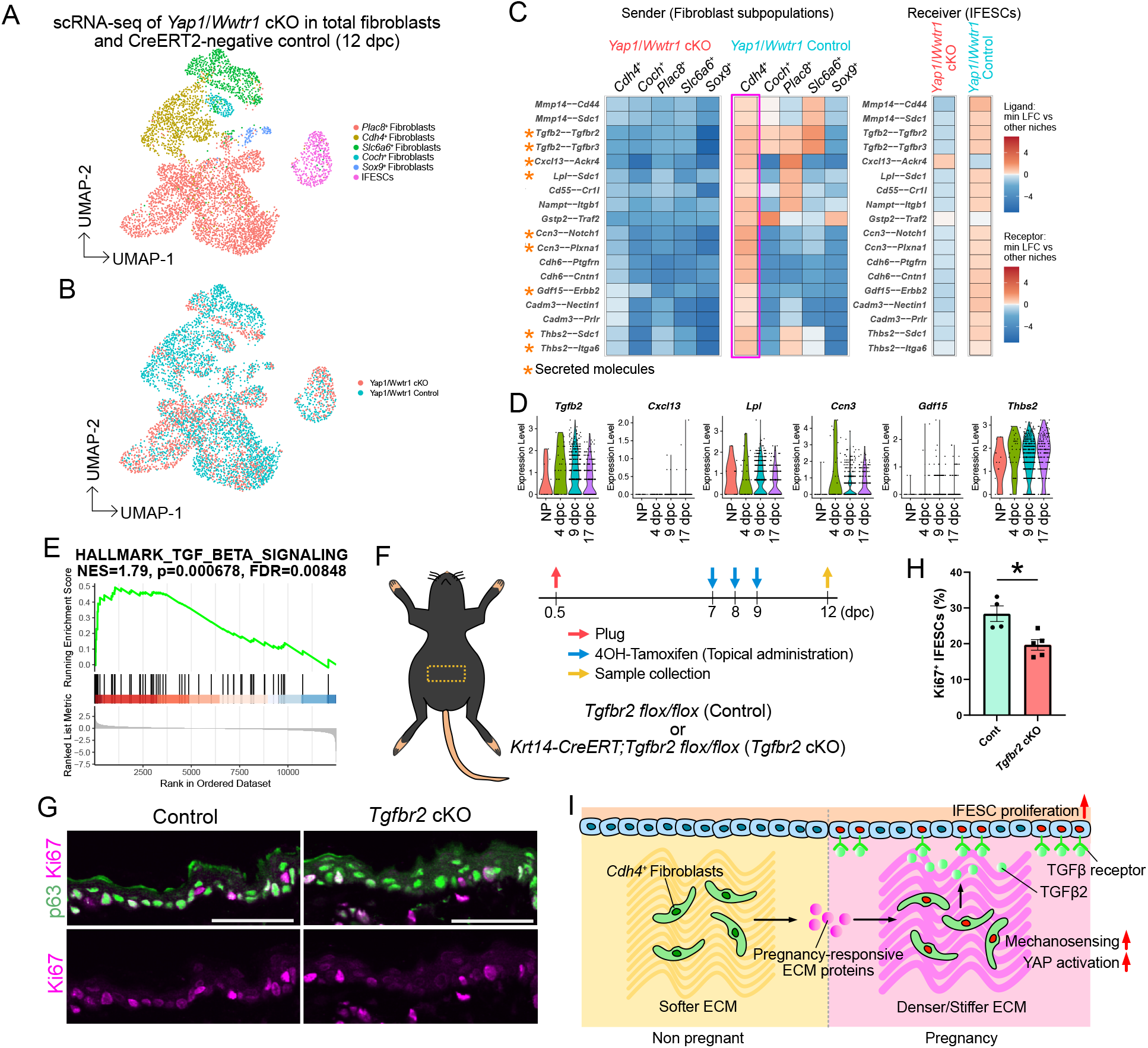
*Cdh4*^+^ fibroblasts induce IFESC proliferation through TGFβ signaling. (**A**) UMAP visualization of five fibroblast subpopulations and IFESCs in scRNA-seq analysis of the *Yap1/Wwtr1* double cKO in total fibroblasts. (**B**) UMAP visualization colored by red and blue that indicate *Yap1/Wwtr1* cKO mice and control, respectively. (**C**) Differential Nichenet analysis. Asterisks indicate genes encoding secreted molecules. (**D**) Violin plots showing the expression of genes encoding secreted molecules listed by (**C**) in *Cdh4*^+^ fibroblasts in non-pregnant and pregnant wild type mice. (**E**) Gene set enrichment analysis (GSEA) plot showing the enrichment of TGFβ signaling in IFESCs at 16 dpc compared to non-pregnant control. (**F**) Schematic showing *Tgfbr2* cKO in *Krt14*-expressing epidermal stem cells during pregnancy. (**G**) Immunostaining for Ki67 (magenta) and p63 (green) in abdominal skin of the *Tgfbr2* cKO in *Krt14*-expressing epidermal stem cells (right) and CreERT2 negative control (left). (**H**) Quantification of (**G**). Data are presented as the mean ± s.e.m. from four-to-five biological replicates. \**P* < 0.05, evaluated with two-tailed unpaired Student’s t-test. (**I**) Summary schematic of this study. Scale bars: 50 μm (**G**).

## Discussion

Our study demonstrates the mechanism by which dermal mechanotransduction regulates epidermal stem cells during healthy skin expansion in pregnancy, centered on a distinct subpopulation of dermal fibroblasts. *Cdh4*^+^ fibroblasts alter the mechano-architecture of the dermis by producing a pregnancy-responsive matrisome. This is followed by mechano-to-chemical conversion via the YAP-TGFβ2 axis in *Cdh4*^+^ fibroblasts. This mechano-to-chemical conversion enables the stromal-epidermal crosstalk to control the skin expansion during pregnancy (Fig. 5I). The adjustment of stromal stiffness is essential for physiological tissue remodeling, and its failure can lead to pathological lesion. For example, mammary glands undergo the healthy remodeling accompanied with reorganization of ECM components during puberty, pregnancy and lactation, changing tissue stiffness and influencing the cell fate decisions of mammary epithelial cells (*26*). Ex vivo studies reveal that the precise control of matrix stiffness is essential for the glandular tissue development (*27, 28*). Conversely, in skin fibrosis, which is a pathological process that induces excessive deposition of ECM and changes the surrounding microenvironment and mechano-field, the stiffening dermal ECM activates signaling including TGFβ and integrin pathways, leading to the pathological alteration of epidermis (*29*). Although the stiffness changes that we detected in the abdominal skin of pregnant mice in this study were only approximately 1.4-fold, this precise, albeit small, control of tissue stiffness could be essential for the healthy tissue remodeling.

Excessive deposition of fibrotic ECM produced by activated fibroblasts (i.e. myofibroblasts) induces skin fibrosis, hypertrophic scarring and keloids (*30*). During pregnancy, several genes encoding pro-fibrotic ECM proteins, including collagen I, lysyl oxidase (LOX), and SPARC (*31*), are upregulated in the *Cdh4*^+^ fibroblasts, whereas fibrillin-1 (FBN1) (*32*) and calreticulin (CALR) (*33*) are downregulated. On the other hand, an anti-fibrotic protein MMP14 (*34*) is upregulated (Fig. 1H). This suggests that ECM remodeling during pregnancy is partially overlaps with, but is distinct from, fibrosis. Recent studies have shown that an inhibition of YAP/TAZ signaling enables scarless wound healing, indicating the YAP activation in dermal fibroblasts promotes scar formation in skin (*35, 36*). YAP is also activated in *Cdh4*^+^ fibroblasts during pregnancy; however, no scar-like phenotype was observed in the abdominal skin of pregnant mice. As YAP is downregulated in dermal fibroblasts in aging skin (*7*), excessive inhibition of YAP is possible to lead to unhealthy tissue remodeling like what observed in aging. Understanding what leads to the scarless remodeling with YAP activation during pregnancy could provide insights into healthy and scarless skin remodeling and repair.

Our integrated transcriptomic analysis identified *Cdh4*^*+*^ fibroblasts are the key sources to produce altered ECM proteins during pregnancy. Skin fibroblasts have been classified into three subpopulations based on their location within the dermis and surface markers: papillary, reticular and hypodermal (*13*). Papillary fibroblasts (CD26^+^/Lrig^+^/Sca1^-^) reside in a superficial location within the dermis, while reticular fibroblasts (Dlk1^+^/Sca1^-^) reside in a deeper location. Hypodermal fibroblasts (CD24^+^/Dlk1^-^/Sca1^+^) are found beneath the dermis. Based on their developmental origin, fibroblasts can also be classified as Engrailed-1 (*En1*)-lineage positive (paraxial mesoderm derived), paired-related homeobox 1 (Prrx1)-lineage positive (lateral plate mesoderm derived), or En1- and Prrx1-lineage negative (*35, 37*). Recent scRNA-seq analysis combined with *in situ* staining reveals the existence of several fibroblast subpopulation in embryonic mouse skin (*38*). In their analysis, they found gene expression of *Plac8* and *Mfap5* in the interstitial layers of dermis/hypodermis at E14.5. The spatial mapping of the fibroblast subpopulations identified in our study indicates that *Plac8*^+^ fibroblasts belong to the hypodermal subpopulation, which is not controversial to the finding in embryonic skin. A part of *Plac8*^+^ fibroblast subpopulation expresses *Procr* (fig. S1D), the gene encoding CD201, which is found as a marker of fascia fibroblasts (*39*), indicating *Plac8*^+^ fibroblast subpopulation includes populations distributed in multiple layers of the hypodermis. On the other hand, the *Cdh4*^+^ fibroblasts includes the papillary, reticular and hypodermal subpopulations. Intriguingly, *Coch*^+^ and *Slc6a6*^+^ fibroblasts are likely associated to hair follicle distribution, and these are distinct from dermal papilla which is located to the bottom of the hair follicles. Identification of functional differences between these subpopulations and correlation to the subpopulations described in previous studies is limited in this study. Further exploration of dermal fibroblast subpopulations is required to provide insight to study dermal cell orchestration.

The skin of mice differs from that of humans in several aspects, including thickness and mechanical properties; Mouse skin has a thinner epidermis and dermis, and exhibits loose skin with lower resting skin tension. Human skin is much thicker, tighter and has higher resting skin tension (*40, 41*). This fact indicates human skin experiences higher tension during pregnancy compared to mice. The stretch of the skin in pregnancy is one of the factors leading to SG in human. SG starts with elastolysis in the dermis, affecting the ECM reorganization with lower expression of collagen and fibronectin, or high proportion of rigid cross-linked collagen (*42*), indicating dysregulation of healthy ECM remodeling can lead to SG. The fact that women with EDS due to TNXB deficiency and other ECM-gene defects rarely develop SG during pregnancy suggests that skin flexibility, which can be achieved by precise ECM remodeling, is essential for the healthy skin expansion during pregnancy. Further investigation into the molecular mechanisms of healthy skin remodeling using a pregnant mouse model could lead to the development of therapeutics for treating or preventing SG and fibrosis.

## Acknowledgements

We thank the members of Toyoshima lab for discussion, Innovative Support Alliance for Life Science at Kyoto University and Stem Cell Laboratory at Institute of Science Tokyo for using shared resources, Single-Cell Genome Information Analysis Core (SignAC) in ASHBi for NGS sequencing, Animal facility of LiMe, Kyoto University and Animal Research Facilities of BioScience Center, Institute of Science Tokyo for mouse experiments. We acknowledge T.Era for providing *Pdgfra-EGFP-CreERT2* mouse line.

## Funding

This research was supported by JST CREST #JPMJCR2023 (F.T., T.Y., M.I., Ko.M.), JST FOREST Program #JPMJFR206C (T.Y.), JSPS KAKENHI #22K15023 (Y.K.), #23K24356 (F.T.), #23K17196 (Ko.M), JSPS A3 Foresight Program #JPJSA3F20230001 (T.Y.), AMED-CREST #JP21gm1310011 (T.Y.), Takeda Science Foundation (F.T.), Naito Foundation (F.T.), KOSE Cosmetology Research Foundation (F.T.), Nanken-Kyoten, Science Tokyo (2023, 2024, 2025), Multilayered Stress Diseases (JPMXP1323015483), Science Tokyo, Medical Research Center Initiative for High Depth Omics, Science Tokyo, ASHBi, supported by the World Premier International Research Center Initiative (WPI) and MEXT Japan.

## Author contributions

Conceptualization: Y.K. and F.T. Methodology: Y.K., K.S., Ko.M., H.F., H.M., A.T., P.R.T., M.I., T.Y. Investigation: Y.K., K.S., Ko.M., H.F., H.M., S.O. M.Y., R.M., M.I., T.Y., F.T. Resources: T.A., Ke.M. Writing: Y.K., K.S., Ko.M., M.I., T.Y., F.T.

## Competing interests

We declare no competing interests associated with this manuscript.

## Supplementary Materials

Materials and Methods

figs. S1 to S7

Table S1

References (43-62)

## Notes

### Competing Interest Statement

The authors have declared no competing interest.

