## Supplemental Figures and Methods for "Distinct dermal fibroblasts direct mechano-chemical signaling to the epidermis during pregnancy"

*Corresponding authors

**The PDF file includes:**

Materials and Methods
Figs. S1 to S7
References

### Materials and Methods

#### Mice

Female mice aged between 8-15 weeks were used for experiments. All of mice were maintained in C57BL/6 strain unless otherwise specified. The mouse lines *B6.Cg-Tnxb*$^{tm1Kmat}$ (*Tnxb* $^{-/-}$)(*43*) and *B6.Cg-Pdgfra*$^{tm1.1(EGFP/cre/ERT2)Hyma}$ (*Pdgfra-CreERT2*)(*44*) were obtained from Riken BioResource Research Center. *Wwtr1*$^{tm1.2Hmc}$ *Yap1*$^{tm1.2Hmc/WranJ}$ (*Yap1 flox;Wwtr1 flox*, maintained as mixed background)(*45*) and *Tg*(*KRT14-cre/ERT*)$^{20Efu/J}$ (*KRT14-CreERT*)(*46*) were purchased from the Jackson Laboratory. *B6.129S6-Tgfbr2*$^{tm1Hlm/Nci}$ (*Tgfbr2 flox*)(*47*) was obtained from the National Cancer Institute Mouse Repository. To get pregnant mice, females were added to male cages individually at 18:00 pm ± 2 h and kept until plug checking at next morning. The pregnant mice that had more than five normal embryos were used for experiments. For *Yap1* and *Wwtr1* conditional knockout (cKO) and *Tgfbr2* cKO, 4-hydroxytamoxifen (4-OHT, Sigma H6278, 1 mg dissolved in 100% ethanol) was topically administrated to shaved abdominal skin at 7, 8 and 9 days post coitum (dpc). The animal experiments were approved by the Committee for Animal Experiments of the Institute for Life and Medical Sciences, Kyoto University and the Institutional Animal Care Committee of Institute of Science Tokyo.

#### Single-cell RNA-seq of mouse skin tissues

Mice were sacrificed by cervical dislocation, followed by harvesting skin tissue. The skin tissue was incubated on Dispase (Corning 354235, 16 U/ml) for 30 min at 25°C, then epidermis was removed by razor blade. The remaining dermal and hypodermal tissues were minced and incubated in enzymatic solution containing Dispase (5 U/ml), Collagenase I (Thermo 17100–017, 450 U/ml) and DNase I (Roche 10104159001, 0.33 U/ml) for 30 min at 37°C with rotation. Dissociated dermal cells were centrifuged at 300*g* at 4°C for 5 min, washed by DMEM/F12-Ham and PBS (-). Dermal cells were incubated with TrypLE at 37°C for 20 min. Cells were filtrated using 70-µm and 35-µm strainers, then resuspended in 0.4% BSA dissolved in PBS (-). Single-cell collection and library preparation were performed using Chromium 3' v3.1 kit (10x Genomics 1000-027) with Chromium Controller or Chromium X. Sequencing was performed using NovaSeq 6000.

#### Bioinformatic analyses

FASTQ files of the output of single-cell RNA-seq of mouse dermal cells were processed using Cell Ranger software v7.1.0 (10x Genomics) using mm10 genome reference. Unique molecular identifier (UMI) counts were further analyzed using R package Seurat v5 (*48*). Background UMI counts were filtered using SoupX package v1.6 (*49*). Cellular multiplets were identified and removed using doubletFinder package v2.0 (*50*). Low quality cells identified the definition of 1,000 < nFeature_RNA < 7,000 and mitochondrial UMI > 15% were removed. UMI counts were normalized using SCTransform (*51*). The number of significant dimensions of principal components were estimated using ElbowPlot function. Cellular clustering was performed based on Shared Nearest Neighbor (SNN) graph. Uniform manifold approximation and projection for dimension reduction (UMAP) rendering was performed to visualize cell clusters. Major cell populations were identified based on the expression pattern of genes shown in fig. S1C. The fibroblast subset was re-clustered, and five subtypes of fibroblasts were identified based on the gene expression pattern shown in Fig. 1C. Fibroblast subset markers based on findAllMarker function were utilized for enrichment analysis using Reactome Pathway database (*52*) on Enrichr (*53*). Genes included in GO:0031012 were utilized to perform principal component analysis (PCA) to generate the matrisome PCA plot. YAP signature gene list was obtained from GSEA "Cordenonsi_YAP_Conserved_Signature", which is originally published in (*54*). To determine the cell-cell interactions between fibroblast subtypes and interfollicular epidermal stem cells (IFESCs), differential NicheNet (v2.2.1) analysis was performed (*55*). For Bulk RNA-seq re-analysis, the RNA-seq profile of abdominal IFESCs obtained from non-pregnant and pregnant mice deposited under GEO accession no. GSE151212. Gene Set Enrichment Analysis (GSEA) was performed using an R package clusterProfiler v4.16 (*56*).

### RNA-HybISS

For RNA-HybISS, harvested tissues were immersed in 4% PFA and incubated at 4°C for 24 hours. Tissues were washed with PBS (-) three times then immersed in 30% sucrose dissolved in PBS (-) at 4°C for overnight. 30% sucrose was replaced with 1:1 solution of O.C.T. compound and 30% sucrose and incubated at 4°C for 2 hours. Tissues were embedded in 100% O.C.T compound and frozen on liquid nitrogen. The tissues were cryo-sectioned at 8-µm thickness and placed on MAS-coated slide glasses (Matsunami, MAS-01). HybISS was performed as described previously (*18*), with modifications that amplicons were produced from padlock probes directly hybridized to RNA species; further methodological details are provided in another study (in preparation). Briefly, slides were post-fixed with 4% formaldehyde for 5 minutes, washed with PBS twice, permeabilized with 0.1 M HCl for 5 minutes and washed with PBS twice. After passing through 70% and 100% ethanol for dehydration, padlock probes were hybridized to RNA, the gaps between 5' and 3' end of probes were ligated, and amplicons were produced with rolling circle amplifications. The resulting amplicons were subjected to sequencing by hybridization. Each round of sequencing included bridge probe hybridization (0.2 µM each) in 1x hybridization buffer (2x SSC, 20% formamide), detection probe hybridization (0.2 µM each) with Hoechst staining (1 µg/mL) in 1x hybridization buffer, imaging, and stripping with stripping solution (2x SSC, 65% formamide). For the visualization of epidermis, immunostaining was performed after RNA-HybISS (fig. S2B). After final probe stripping, slides were blocked with 3% BSA/PBS for 45 min at RT, incubated with anti-pan-Cadherin antibody (Abcam, ab16505, 1:500 dilution) overnight at 4 °C, washed with 0.05% Tween-20/PBS, and incubated with Alexa Fluor 555-conjugated goat anti-rabbit IgG (Thermo, A21428, 1:400 dilution) for 90 min at RT. For the padlock probe design, 1 to 9 target sequences per gene were selected using the Python padlock design software package (https://github.com/Moldia/multi_padlock_design) with the following parameters: arm length, 15 (if targets were not found, 18); Tm, low 65, high 75. Every set of padlock probes for a given gene carries a unique 20 nucleotide (nt) ID sequence. All the sequences of padlock probe targets, bridge probes, and detection probes were shown in supplemental Table 1.

Imaging was performed using a standard epifluorescence microscope (Nikon Ti2-E) connected to an LED light source (Lumencor SPECTRA X light engine). Images were obtained with a CMOS camera (ORCA-Flash4.0V3, Hamamatsu) with a CFI Plan Apochromat Lambda 20x objective (0.75 NA). Filter cubes for wavelength separation were as follows: Chroma 89402X (Hoechst, Cy5), Chroma 89403X (Alexa Fluor 750), Semrock GFP-A-Basic (Alexa Fluor 488), Semrock Cy3-4040C (Cy3), and Semrock CFP-2432C (Atto 425). For multiplex analyses, we assigned a unique color code to each gene, with each code composed of four different fluorophore-conjugated detection probes (Alexa Fluor 750, Alexa Fluor 488, Cy3, and Cy5). Genes were divided into two groups and analyzed in two cycles, each consisting of three rounds of sequencing by hybridization. *Eef1a1* and *Dcn* were imaged separately with Atto 425 to avoid optical crowding. Multispectral images were obtained for multiple cycles. Each image consists of multiple tiles that together cover the tissue section (15% overlap), and each field of view consists of Z-stacks of 1.0 μm steps through the entire tissue thickness. The tiles were stitched together, and the Z-stacks were merged to maximum-intensity projections (NIS-Elements). 16-bit TIFF images were exported and used for subsequent data analysis.

The decoding of gene spots were performed as follows. Firstly, images were coarsely aligned to the reference images (first-round staining images) using a rigid-body algorithm based on Hoechst staining. Then, the images were filtered by subtracting a Gaussian-blurred version of the image ($\sigma$ = 3 pixels) from the original, and composite images were generated from the filtered images of the four detection probe channels. The images were split into multiple tiles, each including a margin, aligned to the reference tiles based on the composite images, and the tiles were stitched back to the original size. Gene spots were detected using the Laplacian of Gaussian filter ($\sigma$ = 1–3 pixels; detection threshold = 0.05) on the first-round image, and the signal intensities in each round were quantified from the filtered images. For each channel, spot intensities were normalized to its 99th percentile value (set to 1). As part of quality control, spots were excluded from the analysis if (i) the maximum intensity among the channels was less than 0.15, or (ii) the ratio of the maximum intensity to the sum of all channel intensities was less than 0.5. Spots that passed through the quality control were used to determine the color code. All the python code used is available upon request from the corresponding author.

**Downstream analysis of RNA-HybISS data**

Nuclear regions were segmented using a Cellpose 2.0 model (*57*) that was custom-trained on our dataset. Nuclear boundaries were expanded by 10 pixels (3.4 μm) outwards using CellProfiler (v4.1.3) (*58*), and the resulting label images representing cells were used for subsequent analyses. Fibroblasts were defined as cells that had more than three spots for *Dcn*. *In situ* cell typing was performed using pciSeq (*19*) with input data including the fibroblast label images, RNA coordinates obtained from RNA-HybISS, and the corresponding scRNAseq data. Cell types and gene expression levels were visualized using napari v0.4.19 (*59*). For cell type visualization and proportion analysis, cells whose maximum probability exceeded 85% were selected.

**Primary mouse dermal fibroblast culture**

Mouse skin tissue digestion was performed as described in 'Single-cell RNA-seq of mouse skin tissues' section. Dermal total cells were resuspended in mouse dermal fibroblast medium (mDFM), a modified medium of previous study (*60*). mDFM is composed of Advanced DMEM/F12 Ham (Thermo, 12634010), Glutamax (Thermo, 35050061), 15 mM HEPES (Thermo, 15630080), 1x B-27 supplement (Thermo, 17504044), 1x N-2 supplement (Thermo, 17502048), 1.25 mM N-acetyl cysteine (Sigma, A9165), mouse Noggin (PeproTech, AF-250-38), mouse PDGFA (Sino Biological, 50447-M07Y), 10 ng/ml mouse FGF2 (Biolegend, 713204) and 1x antibiotic-antimycotic (Thermo, 15240062), and seeded on a well of a six-well plate coated with 5% Matrigel (Corning 354234), cultured at 37°C with 5% $CO_2$. The medium was changed every 2-3 days. The cells at passage 2 were stained with antibodies (FITC-conjugated anti-mouse PDGFRA (eBioscience, 11-1401-82), BV421-conjugated anti-mouse CD31 (Biolegend, 102424), BV421-conjugated anti-mouse CD45 (Biolegend, 103134), Pacific Blue-conjugated anti-human/mouse

CD49f (Biolegend, 313620) eFluor 450-conjugated anti-mouse EpCAM (eBioscience, 48-5791-80). A population of PDGFRA-positive under the negative gating for CD31, CD45, CD49 and EpCAM was collected using FACS sorter MA900 (SONY) or MoFlo XDP (Beckman). Sorted fibroblasts were resuspended in mDFM supplemented with 1 μM mitomycin C cultured on hydrogel on coverglass dish (MatTek) for 24 h. Medium was replaced with ice-cold 4% PFA, and fibroblasts were incubated for 15 min. The fibroblasts were washed using PBS-T, blocked using 1% BSA dissolved in PBS-T at RT for 30 min, incubated with primary antibodies at 4°C overnight.

### Evaluating effects of matrix stiffness on YAP nuclear translocation in cultured dermal fibroblasts

TrueGel3D Hydrogel kit (Sigma, TRUE6) was used to prepare dextran-PEG-based hydrogel. Mixtures containing water, 0.8x TrueGel3D buffer, 3.5 mM (Gel 3.5), 5.0 mM (Gel 5.0), 6.0 mM (Gel 6.0), 7.0 mM (Gel 7.0), or 9.0 mM (Gel 9.0) SLO-dextran and RGD peptide (Sigma, TRUE-RGD) were incubated for 20 min at RT, followed by addition of 20% mDFM and 2.8 mM (Gel 3.5), 4.0 mM (Gel 5.0), 4.8 mM (Gel 6.0), 5.6 mM (Gel 7.0), or 7.2 mM (Gel 9.0) PEG non-cell-degradable crosslinker. The mixtures were placed on glass bottom dishes (MatTek, P35G-0-14-C) and incubated at 37°C for 70 min. For stiffness measurement by AFM, the solidified gels were immersed in PBS until the measurement. For testing YAP1 nuclear translocation, FACS-sorted primary mouse abdominal dermal fibroblasts (40,000 cells per 200 μl mDFM) are seeded on the solidified hydrogel and incubated at 37°C for 2 hours. We added 3 mL of mDFM supplemented with Mitomycin C (Nacalai Tesque, 20898-21) was added to the dishes then incubated for additional 22 hours. The fibroblasts were fixed using 4% PFA in PBS. For blocking, the slides were incubated with 1% bovine serum albumin (BSA, Wako, 018-15154) dissolved in PBS supplemented with 0.1% Triton X-100 (PBS-T) at RT for 30 min, followed by incubation with YAP1 antibody (Cell Signaling, 4912S, 1:400 dilution) diluted in 1% BSA at 4°C overnight. Primary antibody was washed by PBS-T, followed by incubation with Cy3-conjugated donkey anti-rabbit IgG secondary antibody (Jackson ImmunoResearch, 711-166-152, 1:400 dilution) at RT for 45 min. The coverslips were removed from dishes using Coverslip Removal Fluid (MatTek, DCF-OS-30), then mounted on slideglasses using Fluoromount-G mounting medium with DAPI (Thermo, 00-4959-52). Imaging was performed using confocal microscope FV3000 (Olympus) using 20x objectives. YAP1-positive nuclei were determined as > 20% YAP1-positive area in DAPI-positive nuclei area analyzed using celSense software (Olympus).

### Atomic Force Microscopy (AFM)

Mouse skin tissues were harvested and immediately embedded in O.C.T. compound on liquid nitrogen. The tissues were cryo-sectioned at 8-μm thickness and placed on MAS-coated slide glasses. The slides were washed with PBS (-) three times, stained with Hoechst 33342. For the topographical measurement, the cryo-sectioned tissues were fixed with 4% PFA for 15 min. Hydrogels were placed on the slide glasses. These samples were mounted on atomic force microscope system (JPK BioAFM NanoWizard 3; Bruker Nano GmbH., IX81; Olympus Co.) and examined by the Quantitative Imaging™ mode, which measures the surface topography and the stiffness of the sample. AFM cantilevers (qp-BioAC CB2; Nanoworld AG) were calibrated with a thermal noise method (*61*). For the skin tissue, the dermal regions were identified based on Hoechst staining patterns. The set point was 300 pN for the stiffness measurement of the non-fixed tissues and hydrogels, and 500 pN for the topographical measurement of the fixed tissues. The piezo displacement speed was 50 μm/s in all measurements. Based on the force ($F$) versus indentation depth ($h$) curves obtained in the Quantitative Imaging$^{TM}$, slope (nN/μm) was estimated with a linear regression for the sample points within the force range of 100 pN ≤ $F$ ≤ 300 pN.

#### DNA-based digital tension sensor

Both single-stranded DNA (ssDNA) oligonucleotides comprising the tension sensor were synthesized by Integrated DNA Technologies. One oligonucleotide was modified with a biotin group at the 5'end and a thiol group at the 3'end. The complementary oligonucleotide was modified with a cholesterol group at the 5'end and a Cy3 fluorophore at the 3'end. Cyclic RGDfK (cRGDfK; Biosynth, PCI-3696-PI) was conjugated to the thiol-modified oligonucleotide as follows. cRGDfK (8 mM) was mixed with sulfo-SMCC (4 mM; Thermo Scientific, A39268) and incubated for 20 min at RT. The resulting cRGDfK-SMCC conjugate was reacted with 400 μM thiol-modified oligonucleotide and incubated overnight at 4°C. Labeling efficiency of cRGDfK to the thiol-modified oligonucleotide was evaluated by electrophoresis on a 20% TBE polyacrylamide gel (Life Technologies) and determined to be >95%.

To assemble the tension sensor, 6 μM cRGDfK- and biotin-modified oligonucleotides were mixed with 5 μM cholesterol- and Cy3-modified complementary oligonucleotides in TE buffer. The mixture was annealed by heating at 90°C for 5 min, followed by cooling at a rate of 3.5°C per minute. The assembled sensor was aliquoted and stored at −80°C until use.

#### Immobilization of the tension sensor onto hydrogels

Hydrogels were prepared using the TrueGel3D Hydrogel Kit (Sigma-Aldrich, TRUE6). Mixtures containing water, 0.8x TrueGel3D buffer, 5 mM (for Gel5.0) or 7 mM (for Gel7.0) SLO-Dextran, and 0.533 mM SH-PEG-Biotin (Biopharma PEG Scientific, HE003041-1K) were incubated for 1 hour at RT. These were then mixed with 20% DMEM/F-12 Ham (Nacalai Tesque, 08460-95) and 4 mM (Gel5.0) or 5.6 mM (Gel7.0) PEG non-cell-degradable crosslinker. These mixtures were immediately spread into the hole of glass-bottom dishes (Phoenix Science, P35G-0-14-C/H) and incubated for at least 1 hour at 37°C to form hydrogels. The hydrogels were washed three times with PBS and incubated with 2.5 mg/ml NeutrAvidin (Invitrogen, A2666) for 1 hour at 37°C. After three PBS washes, the hydrogels were incubated with 500 nM 30 pN-tension sensor for 1 hour at 37°C, followed by three additional PBS washes.

#### Preparation and seeding of mouse dermal fibroblasts on tension sensor hydrogels

Mouse dermal fibroblasts, isolated and cryopreserved at -150 °C, were thawed immediately before use and cultured in mDFM with Matrigel under the conditions described above, without further passaging. Mouse fibroblasts (4–7 × $10^4$ cells/mL) were seeded onto hydrogels functionalized with the tension sensor and incubated in mDFM at 37°C under 5% $CO_2$. For the 6-hour incubation, 1 μM mitomycin C (Nacalai Tesque, 20898-21) in mDFM was added to the glass-bottom dishes 2 hours after cell seeding. After 6 hours of incubation, cells were gently rinsed three times with PBS and treated with TrypLE Express (Gibco, 12604021) for 5 minutes at 37°C, followed by gentle pipetting to detach them from the tension sensor. Detached cells were then reseeded onto multi-well glass-bottom dishes (Matsunami Glass, D141400) coated with 50 mg/mL BSA (Sigma-Aldrich, A2153). DMEM/F-12 Ham supplemented with 10% FBS (Gibco, A3840002) and 1x Antibiotic-Antimycotic was added to inhibit enzymatic activity.

#### 3D imaging and analysis of the Cy3 tension sensor attached to cells

Fluorescence imaging of the Cy3 tension sensor attached to cells was performed using an Evident IX83 microscope equipped with a multiline solid-state 555 nm laser for Cy3 fluorescence (89 North, LDI-7), a short-arc lamp for bright-field imaging (Evident, U-HGLGPS), a 60x objective lens (Evident, PlanApo N 60x), a confocal scanner unit (YOKOGAWA, CSU-W1), dichroic mirrors (405/488/561/640 nm), band-pass filters (617/73 nm), and a digital CMOS camera (Hamamatsu, ORCA-Flash4.0), all controlled using cellSens software (Ver. 2.3, Evident). Z-stack images were acquired by shifting the focal plane in 1-µm increments from the bottom to the top of the cell, with each z-slice captured at 2 Hz for Cy3 imaging (Fig. S5).

Cells with too low fluorescence of Cy3 to be distinguished from background or with ruptured membranes were excluded from analysis. Image processing was performed using ImageJ (Ver. 1.54p). Each z-slice was processed with a median filter and binarized to detect cell shape. Although Cy3 fluorescence was observed along the entire cell membrane, its intensity was non-uniform. When necessary to facilitate analysis, the cell membrane was manually traced, and fluorescence outside the membrane was removed prior to binarization. The cell region in each z-slice was identified from the binary image after applying the "Fill Holes" function in ImageJ to measure cell area ($A_k$). The Cy3 fluorescence intensity ($FI_k$) was then measured from the corresponding Cy3 image within the same region (fig. S5). The integrated Cy3 intensity per cell was calculated as $\sum_k(FI_k / A_k)$. The results were confirmed across three independent biological replicates.

#### Immunofluorescence

Mouse skin tissues were harvested and immediately embedded in 100% OCT compound and frozen on liquid nitrogen. The tissues were cryo-sectioned at 8-µm thickness and placed on MAS-coated slide glasses. The slides were immersed in ice-cold 4% PFA for 15 min and were washed using PBS-T. For blocking, the slides were incubated with 1% bovine serum albumin (BSA, Wako, 018-15154) dissolved in PBS-T at RT for 30 min, followed by incubation with primary antibodies (anti-YAP1 (Cell Signaling, 4912S, 1:200 dilution), anti-PDGFRA (R&D, AF1062, 1:200 dilution), anti-Ki67 (eBioscience, 14-5698-82, 1:200 dilution), anti-p63 (GeneTex, GTX124660, 1:200 dilution) or anti-TNXB (M23, homemade (*62*), 1:500 dilution) diluted in 1% BSA or Can Get Signal Immunostain Immunoreaction Enhancer Solution (TOYOBO, NKB-401) at 4℃ overnight. Primary antibodies were washed by PBS-T, followed by incubation with secondary antibodies (Cy3-conjugated donkey anti-rabbit IgG (Jackson ImmunoResearch, 711-166-152), Alexa Fluor 594-conjugated donkey anti-rabbit IgG (Jackson ImmunoResearch, 711-586-152), Alexa Fluor 488-conugated donkey anti-rabbit IgG (Jackson ImmunoResearch, 711-546-152), Alexa Fluor 647-conugated donkey anti-goat IgG (Jackson ImmunoResearch, 705-606-147), Alexa Fluor 488-conugated donkey anti-goat IgG (Jackson ImmunoResearch, 705-546-147), or Alexa Fluor 647-conjugated donkey anti-rat IgG (Jackson ImmunoResearch, 712-606-153) diluted in PBS-T (1:400 dilution) at RT for 45 min. The slides were mounted using Fluoromount-G mounting medium with DAPI (Thermo, 00-4959-52). Imaging was performed using confocal microscope FV3000 (Olympus) using 20x objectives. Proportion of proliferating IFESCs were determined using cellSense (Olympus) and ImageJ software. YAP1 relative fluorescence unit (RFU) in nuclei of dermal fibroblasts was calculated using cellSense.

### Statistical analysis

A two-tailed unpaired or paired t-test was used to compare two groups. An one-way analysis of variance (ANOVA) was followed by Tukey's and Dunnet's tests for multiple comparisons to compare more than two groups. Statistical significance was considered when the *P* values were less than 0.05. All statistical analyses were performed using GraphPad Prism v9 or v10.

fig. S1

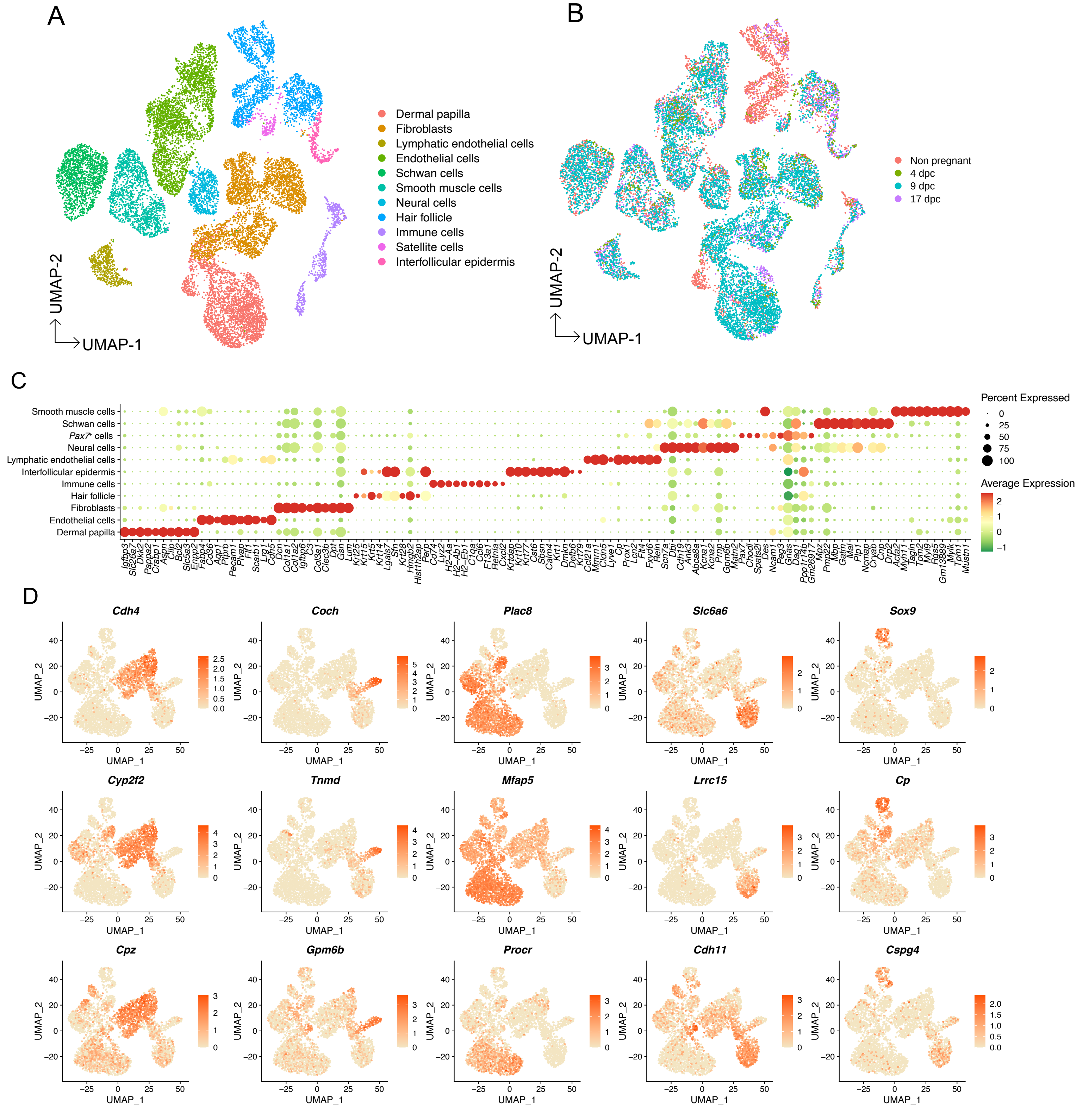


**fig. S1. Single-cell RNA-seq analysis of abdominal skin during pregnancy.** (**A**) UMAP visualization of total cells captured by scRNA-seq analysis. (**B**) UMAP visualization colored by timepoints. (**C**) Dot plot showing cell type markers utilized for distinguishing cell types. (**D**) UMAP showing fibroblasts with expression of their subpopulation markers.

fig. S2

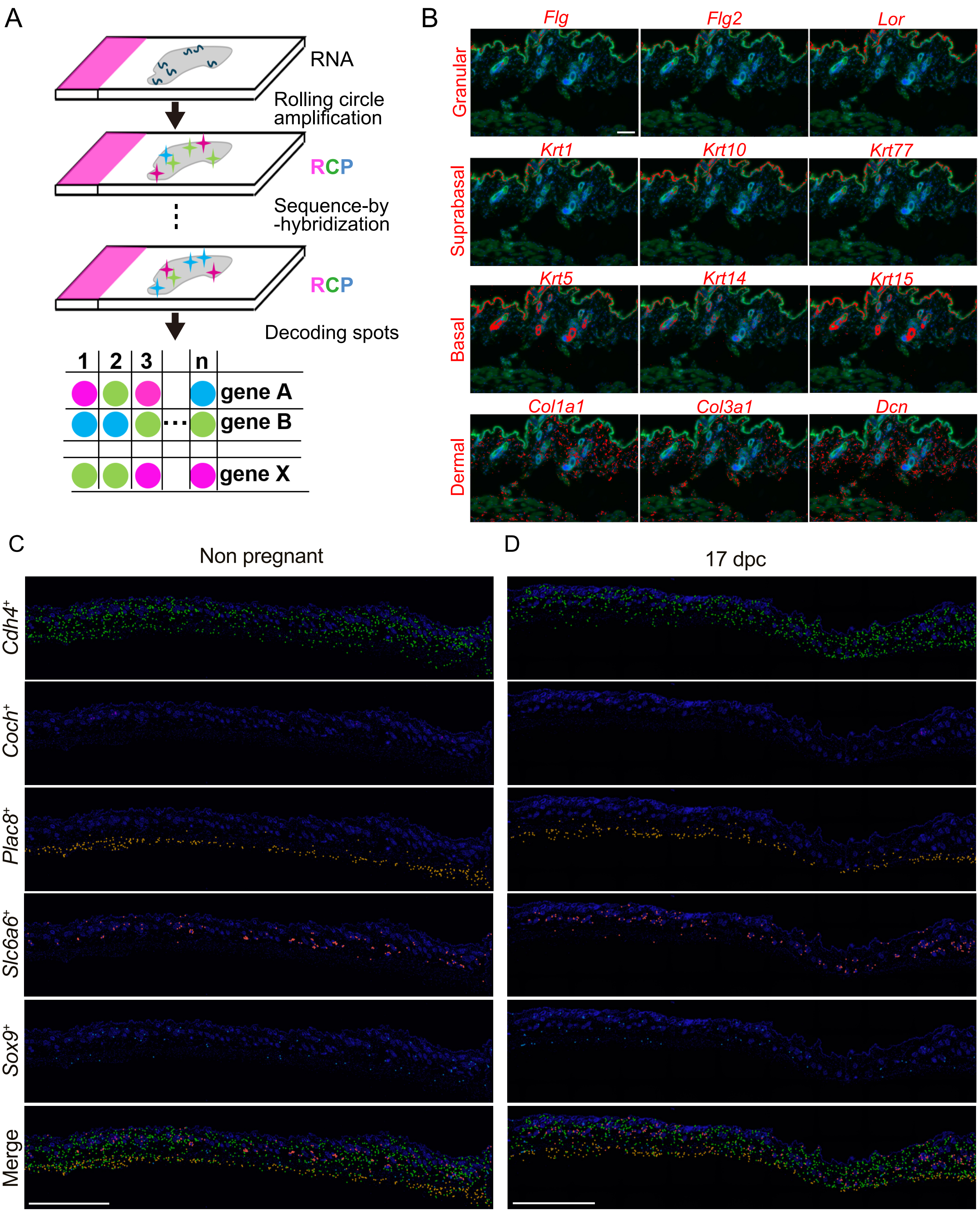


**fig. S2. Optimization for RNA Hybridization-based in situ sequencing (RNA-HybISS) and pci-Seq.** (**A**) Schematic of RNA-HybISS. RCP, rolling circle product. (**B**) RNA HybISS images of abdominal skin. Red, RNA amplicons of indicated genes; Blue, Hoechst 33342; Green, pan-Cadherin antibody staining. (**C, D**) Five subpopulations of fibroblasts determined by pci-Seq in abdominal skin obtained from non-pregnant (**C**) and pregnant (**D**, 17 dpc) mice. Scale bars: 100 μm (B), 1 mm (**C** and **D**).

fig. S3

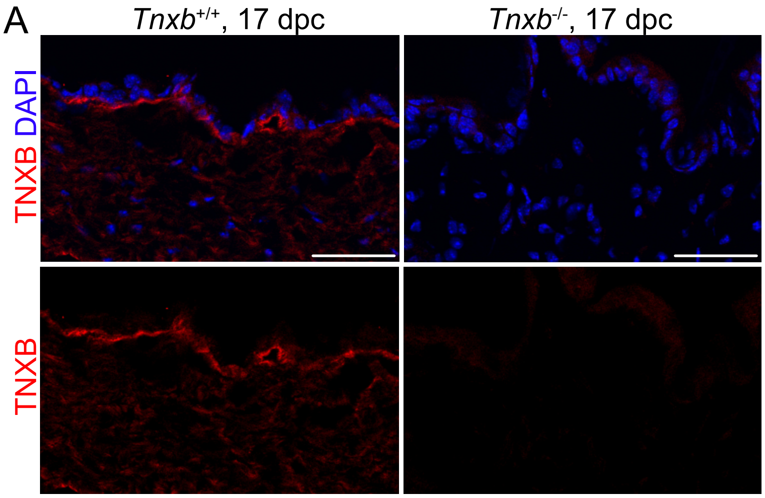


**fig. S3. TNXB staining. (A)** Immunostaining for TNXB (red) of abdominal skin obtained from *Tnxb* KO (right) and control mice (left). Scale bars: 50 µm (**A**).

fig. S4

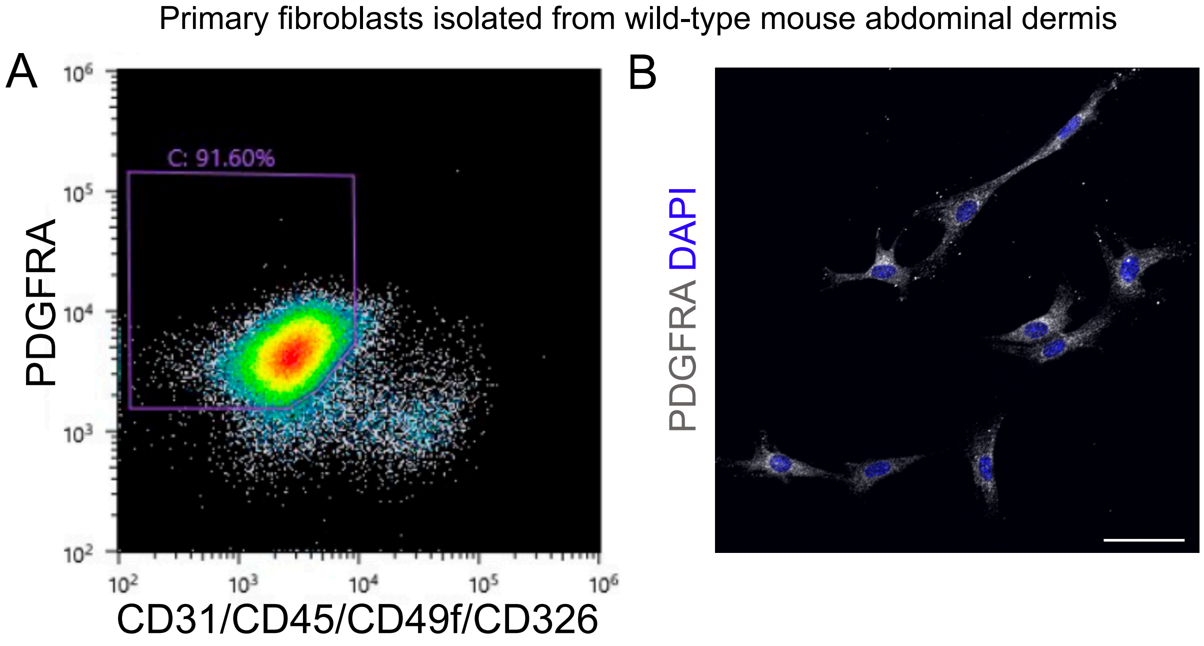


**fig. S4. Primary dermal fibroblast isolation and culture.** (**A**) A FACS plot showing total dermal cells cultured using fibroblast growth medium at passage 2. (**B**) Immunostaining for PDGFRA (gray) in the FACS-sorted primary dermal fibroblasts. Scale bar: 50 µm (**B**).

fig. S5

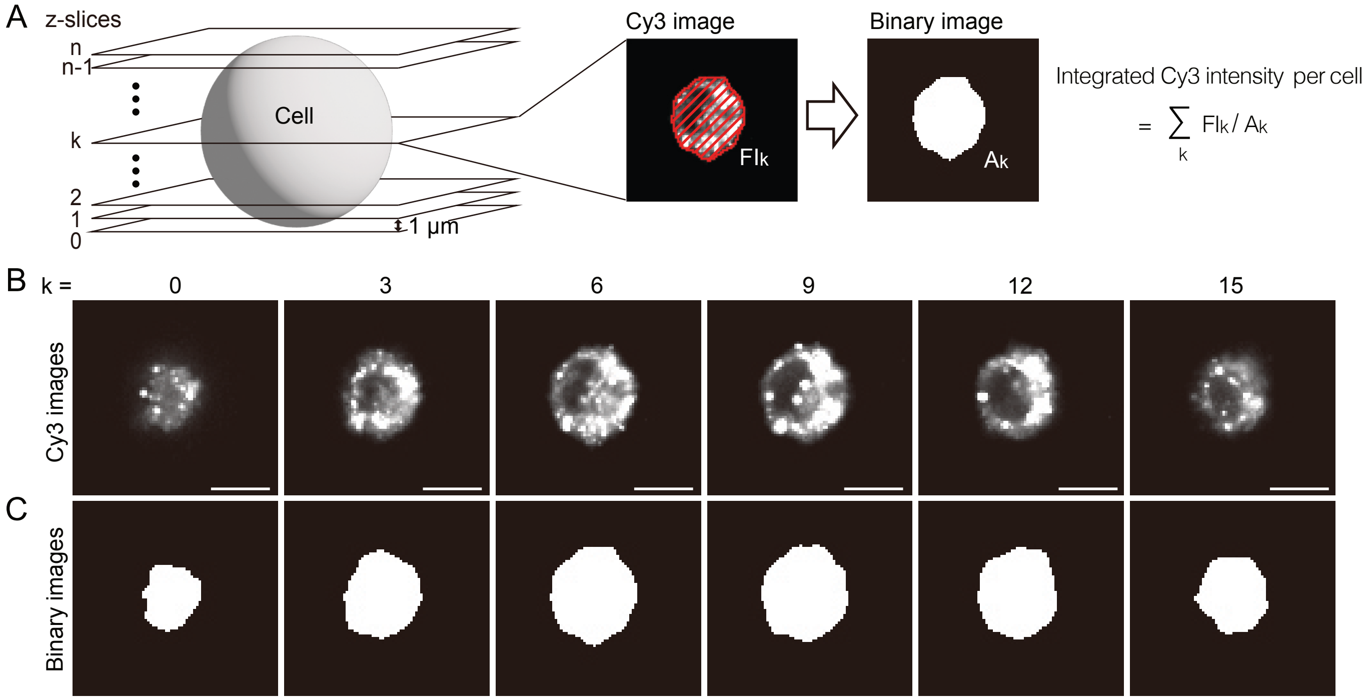


**fig. S5. Measurement of integrated Cy3 intensity per cell.** (**A**) Schematic of z-slices acquired for 3D imaging of a cell. Z-slices were acquired in 1-μm increments from the bottom to the top of the cell. Integrated Cy3 intensity in individual cells was calculated from Cy3 fluorescence intensities (FI$_k$) and areas (A$_k$) measured in each z-slice. The Cy3 image and binary image correspond to those in (**B**) and (**C**), respectively, with k = 6. (**B, C**) Z-slice images of a cell incubated for 1 hour on Gel7.0 coated with 500 nM tension sensor. (**B**) Cy3 images and (**C**) corresponding binary images of the z-slices. Scale bars: 10 μm (**B**).

fig. S6

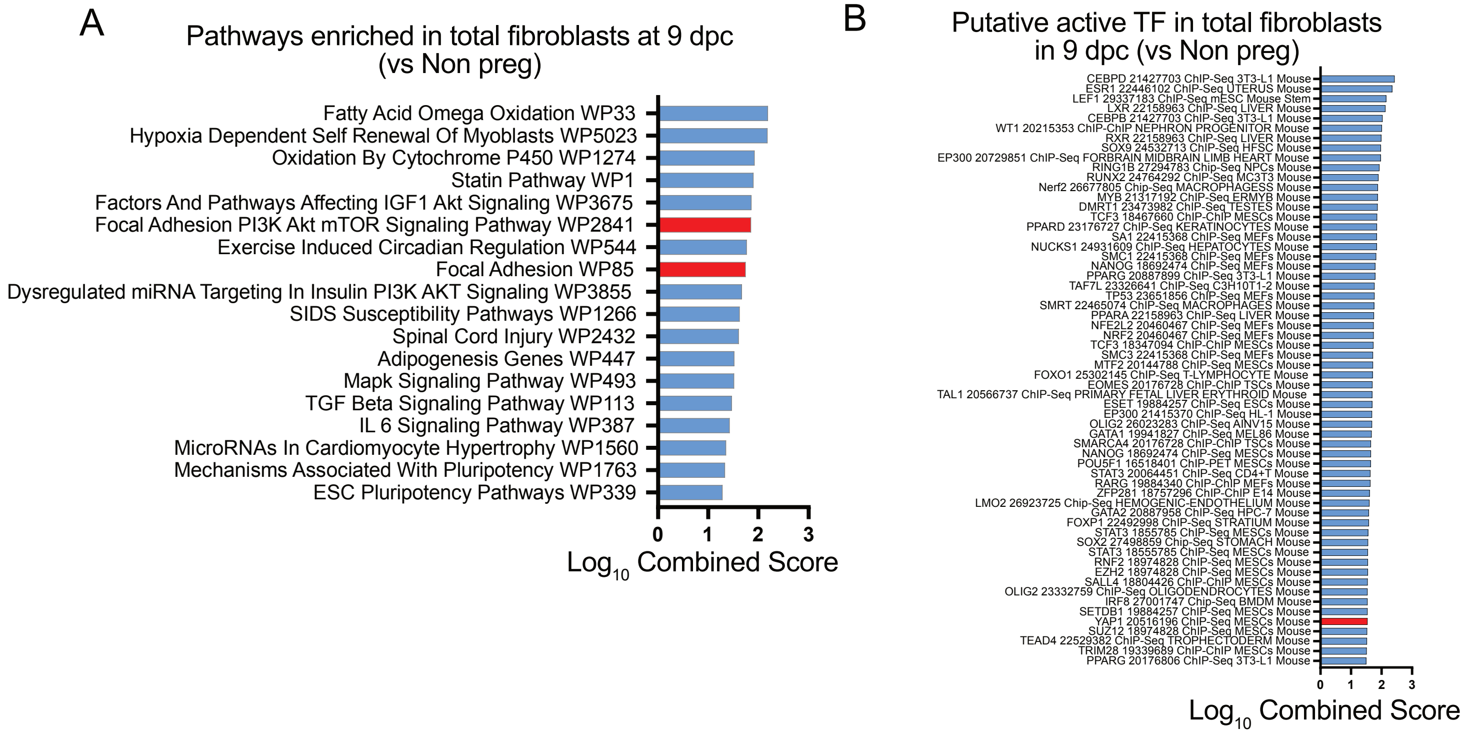


**fig. S6. Mechanosignaling detected in single-cell RNA-seq of mouse abdominal dermal fibroblasts.** (**A**) Enriched pathways in total fibroblasts of scRNA-seq results at 9 dpc compared to non-pregnant control. (**B**) Putative active transcriptional factors (TFs) in total fibroblasts at 9 dpc compared to non-pregnant control.

fig. S7

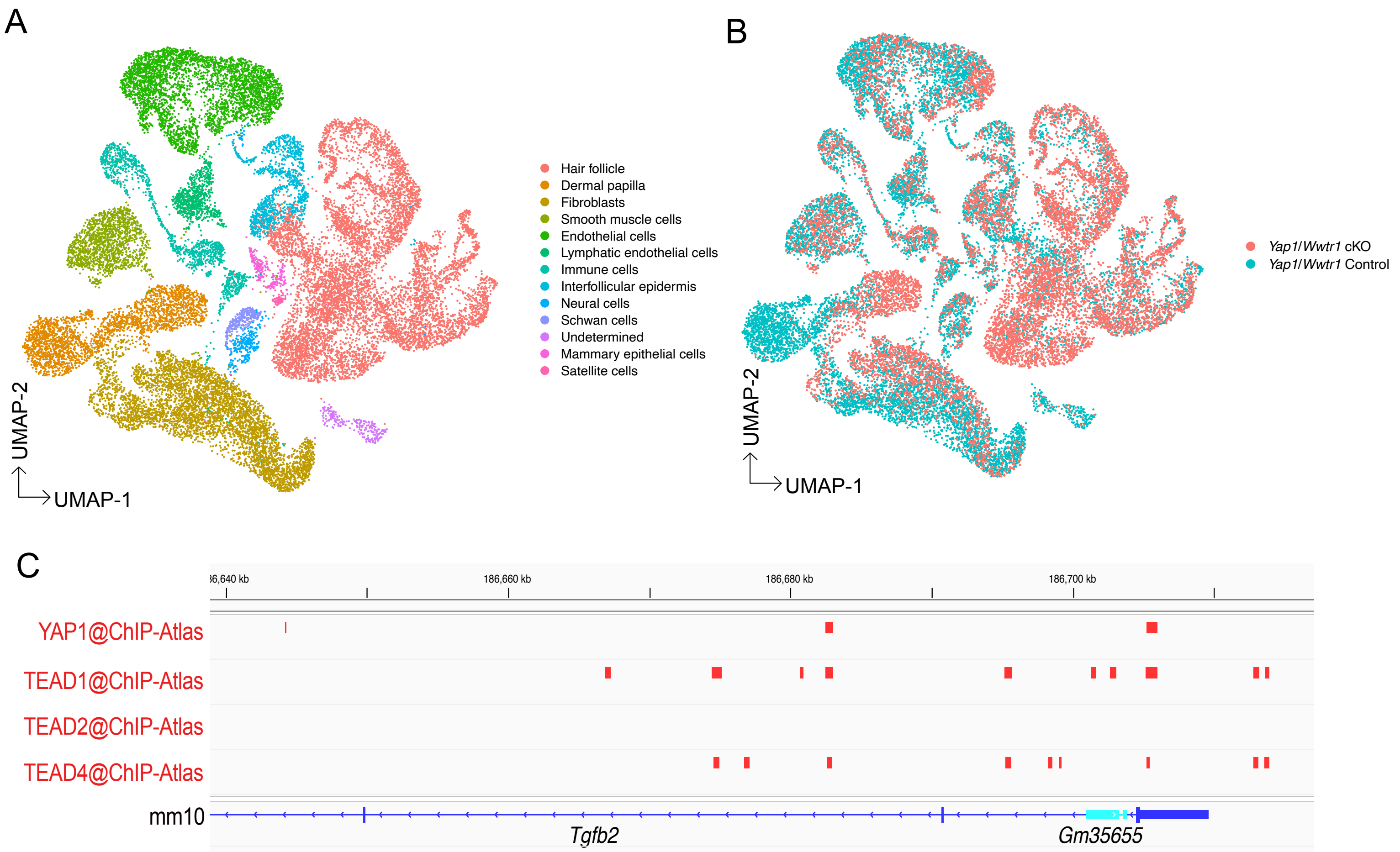


**fig. S7. YAP1 function in skin during pregnancy.** (**A**) UMAP visualization showing total cells obtained from *Yap1/Wwtr1* cKO in total fibroblasts, and control mice at 12 dpc. (**B**) UMAP visualization colored by the sample conditions. (**C**) ChIP-atlas of YAP1, TEAD1, TEAD2 and TEAD4 in the *Tgfb2* locus.
